# Accurate prediction of in vivo protein abundances by coupling constraint-based modelling and machine learning

**DOI:** 10.1101/2023.04.11.536445

**Authors:** Maurício Alexander de Moura Ferreira, Philipp Wendering, Marius Arend, Wendel Batista da Silveira, Zoran Nikoloski

## Abstract

Quantification of how different environmental cues affect protein allocation can provide important insights for understanding cell physiology. While absolute quantification of proteins can be obtained by resource-intensive mass-spectrometry-based technologies, prediction of protein abundances offers another way to obtain insights into protein allocation. Here we present CAMEL, a framework that couples constraint-based modelling with machine learning to predict protein abundance for any environmental condition. This is achieved by building machine learning models that leverage static features, derived from protein sequences, and condition-dependent features predicted from protein-constrained metabolic models. Our findings demonstrate that CAMEL results in excellent prediction of protein allocation in *E. coli* (average Pearson correlation of at least 0.9), and moderate performance in *S. cerevisiae* (average Pearson correlation of at least 0.5). Therefore, CAMEL outperformed contending approaches without using molecular read-outs from unseen conditions and provides a valuable tool for using protein allocation in biotechnological applications.

## Introduction

Proteomics technologies facilitate system-wide profiling of the identity as well as changes in abundance, distribution, modifications, and interactions of proteins that drive different cellular processes (Schubert et al., 2017). As cells are dynamic systems, these characteristics of the proteome change in response to different environmental cues to facilitate the system’s adaptation to different physiological states (Liu et al., 2019; Nielsen, 2019). As a result, high-throughput quantitative proteomics technologies, based on mass spectrometry (MS) techniques, have provided valuable information for various biotechnological and medical applications (Kültz, 2020; Lill et al., 2021). However, the measurement of absolute protein abundance still poses great challenges, due to physicochemical properties that interfere with ionization efficiency in mass spectrometry (Otto et al., 2014; Pappireddi et al., 2019) and the lack of a standardized approach, affecting the reproducibility of resulting data (Calderón-Celis et al., 2018).

Another approach for protein quantification relies on predicting protein abundances by using a variety of machine learning approaches using features that are more facile to measure or quantify. For instance, with gene expression data as predictors, Torres-García et al. (2009) trained gradient-boosted trees to predict protein abundances in *Desulfovibrio vulgaris*, resulting in prediction accuracies, quantified by the coefficient of determination, R^2^, ranging from 0.39 to 0.58. Similarly, Mehdi et al. (2014) used Bayesian networks to predict protein abundance for *Saccharomyces cerevisiae* and *Schizosaccharomyces pombe* using gene expression data, yielding Pearson’s correlation coefficients ranging from 0.50 to 0.71. Mergner et al. (2020) used stepwise and LASSO regression models to predict protein abundances in *Arabidopsis thaliana* based on gene expression as well as sequence-derived features, achieving Pearson correlation in the range from 0.61 to 0.79 across different tissues. Codon usage metrics have also been used to predict protein abundance in *Saccharomyces cerevisiae*, with the best model achieving an R^2^ value of 0.74 (Ferreira et al., 2021). In addition, structural features of mRNA molecules have been used to predict protein abundances in *Escherichia coli*, achieving a Spearman correlation coefficient of 0.71. However, these machine learning models have been developed using quantitative proteomics and transcriptomics data from optimal growth conditions and their performance in sub-optimal growth conditions remains unexplored. In addition, given the dynamic nature of the proteome, it is expected that the predictions across different conditions from machine learning models based on environment-invariant features (e.g., codon usage and structure-related), are poor.

The advent of protein-constrained genome-scale metabolic models (pcGEMs) (Beg et al., 2007) has facilitated not only the prediction of protein abundance but also the usage of proteomics data to predict metabolic and physiological traits (Bekiaris and Klamt, 2020; Domenzain et al., 2022; Sánchez et al., 2017). This is achieved by parameterizing pcGEMs with data on enzyme turnover numbers, providing the basis to link steady-state fluxes with enzyme abundances. For instance, a pcGEM of *E. coli* was used by Adadi et al. (2012) to predict protein abundance, achieving Pearson correlation coefficient of 0.84 with gene expression data of *E. coli* grown on a glucose minimal medium. However, given that comparisons were made with gene expression data, it remains unexplored how these predictions fare against measured proteomics data. Further, Heckmann et al. (2018) trained machine learning approaches to predict enzyme catalytic efficiency, namely *in vitro* (*k*_*cat*_) and *in vivo* values 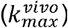, that were later used to predict protein abundances with the approach of Adadi et al. (2012). Comparing the predicted protein abundances to experimental data from Schmidt et al. (2016), they found that using 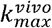 values resulted in a 43% lower root mean squared error (RMSE) in comparison to *k*_*cat*_ values. However, when comparing the prediction to measured proteomics data, the obtained R^2^ values were poor to modest, ranging from 0.11 to 0.68. As a result, we still lack an understanding of the factors that affect the quality of the protein abundance predictions. Moreover, like the machine learning approaches above, the existing predictions of protein abundance based on pcGEMs have only been assessed under conditions that achieve maximal measured specific growth rate for the wild-type strain that we refer to as optimal. Therefore, the problem of predicting protein abundance under sub-optimal, stress-related conditions, that result in smaller specific growth rate than the optimal, is particularly difficult since many mechanisms affecting protein abundance (e.g., codon usage (Novoa et al., 2019), RNA levels (Eraslan et al., 2019), protein interactions (Mergner et al., 2020), 3’ or 5’ UTR motifs (Terai and Asai, 2020)) are not included in pcGEMs.

To address these issues, we coupled constraint-based modelling with machine learning to predict *in vivo* protein abundances, leading to a multi-model framework termed CAMEL (Coupled Approach of MEtabolic modelling and machine Learning) that leverages the strengths of both approaches. First, CAMEL predicts the protein abundance in different conditions using a constraint-based approach. Next, these protein abundance predictions are employed together with experimentally measured protein abundances to calculate protein reserve ratios. CAMEL then trains machine learning (ML) models to predict the protein reserve ratios using different static features, derived from the protein sequence, and condition-dependent features, obtained from the pcGEM. Lastly, the predicted protein reserve ratios along with the predicted protein abundances from the pcGEM are used to calculate *in vivo* protein abundances. Once trained, the models of CAMEL can be used to make predictions by using features predicted by the pcGEM instead of data on molecular read-outs from unseen conditions. Our results showed that the CAMEL approach outperformed contending methods on data from two model organisms, *E. coli* and *S. cerevisiae*, highlighting the advantages of a coupling of constraint-based modelling and machine learning approaches. In addition, these applications of CAMEL point to particular proteins and metabolic pathways for which protein abundance is difficult to predict, requiring the consideration of additional features to improve prediction performance in future developments.

## Material and Methods

### Data set construction

We obtained the experimental enzyme abundances for *E. coli* over 31 growth conditions from three different studies (Peebo et al., 2015; Schmidt et al., 2016; Valgepea et al., 2013), as reported in Davidi et al. (2016). The growth conditions in these experiments range from alternative carbon substrates (e.g., acetate, glucosamine, glucose, glycerol, mannose, pyruvate, and xylose) to chemostats with increasing dilution rates, with glucose as a carbon source. For *S. cerevisiae*, we used the enzyme abundance values from the data set of Chen and Nielsen (2021), which contains quantitative proteomics measurements for 30 growth conditions obtained from four studies (Chen and Nielsen, 2021; Di Bartolomeo et al., 2020; Lahtvee et al., 2017; Yu et al., 2020), including: optimal batch growth with glucose as carbon source, nitrogen- and glucose-limited chemostats with increasing dilution rates, stress states achieved with high ethanol and osmolarity, and reduced nutrient availability.

Condition-dependent features are predicted from the pcGEMs and include the enzyme usage and the metabolic fluxes associated with each enzyme *i*. These features are obtained by solving a linear problem to minimize excess enzyme usage (Eqs. (1)-(8)), detailed below. To predict the enzyme usage distributions, we used the pcGEMs eciML1515 (Domenzain et al., 2022) and ecYeast8 (Lu et al., 2019), from *E. coli* and *S. cerevisiae*, respectively, both without integrated enzyme measurements (batch model). To this end, we minimized excess enzyme usage. Maximal enzyme usage is linked to efficient usage of resources, which has been shown to result in better estimates of *k*_*cat*_ values over other approaches that focus on minimization of total flux (Xu et al., 2021). This objective function also ensures that cellular burden is alleviated, which can lead to proteotoxic effects for the cell (Kintaka et al., 2020). We solve the following linear programming (LP) problem, termed LP1:

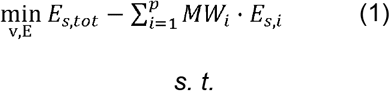

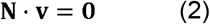

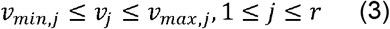

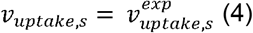

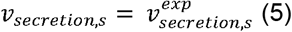

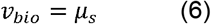

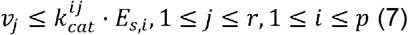

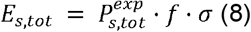

where *E*_*s,i*_ is the enzyme *i*, ranging from 1 to the number of proteins *p* in the model (1 *≤ i ≤ p*), in condition *s, MW*_*i*_ is the molecular weight of an enzyme *i*, **N** is the stoichiometric matrix, **v** is the flux distribution vector, *v*_*j*_ is the metabolic flux through reaction *j*, ranging from 1 to the number of reactions *r* in the model (1 *≤ j ≤ r*), *v*_*bio*_ is flux through the biomass reaction, and *μ* is the specific growth rate, *E*_*tot*_ is the total enzyme content, 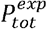 is the total protein content, *f* is the mass fraction of all measured proteins included in the model, and a is the average *in vivo* enzyme saturation, assumed to be a specific value of 0.5 for conditions in which this parameter is not available (Domenzain et al., 2022; Sánchez et al., 2017). The constraints applied to the problem are based on the available data about specific growth rates, secretion, and nutrient uptake rates, protein usage and total protein content.

For both organisms, we constrained the models given the measurements of nutrient uptake rates and specific growth rates using data from Davidi et al. (2016) and Chen and Nielsen (2021). We excluded the experimental conditions that lacked physiological data (e.g., nutrient uptake rates, specific growth rates, or protein content). In addition, we opted to exclude the temperature stress conditions from Lahtvee et al. (2017), as temperature stress triggers responses that entail cellular changes that can impact the function of enzymes (Li et al., 2021) and require more tailored modelling approaches (Wendering et al., 2023). This filtering resulted in data from 20 conditions for *E. coli*, and 19 conditions for *S. cerevisiae*. For conditions for which the models were overconstrained, we relaxed the constraints on the uptake and secretion rates by increments of 1% until the measured specific growth rate could be achieved.

We were also interested in whether using pcGEMs integrated with *in vivo* catalytic rates 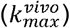 instead of *in vitro* values (*k*_*cat*_) would impact the performance of the ML models. To this end, we exchanged the integrated *k*_*cat*_ values with the 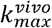 values, obtained from the compilation by (Wendering et al., 2023) of estimated catalytic rates using pFBA from the Davidi et al. (2016) and Chen and Nielsen (2021) studies. For both *E. coli* and *S. cerevisiae*, we concatenated the predictions generated from the considered growth conditions into a single data set for each organism. The resulting data set for *E. coli* included 2256 enzymes for the pcGEM using *k*_*cat*_ values, and a data set with 2246 enzymes for the pcGEM 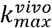 values. For *S. cerevisiae*, the resulting data set included 4596 enzymes for the pcGEM using *k*_*cat*_ values, and a data set with 4590 enzymes for the pcGEM 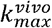 values.

### Training and assessment of machine learning models

To train the machine learning models, we used the Tree-based Pipeline Optimization Tool (TPOT), an automated machine learning tool that optimizes machine learning pipelines to predict the target variable using genetic programming (Le et al., 2020). We configured TPOT for regression problems since we aimed to obtain quantitative predictions of the protein reserve ratio, *ϕ*_*i*_, for each enzyme *i*, 1 *≤ i ≤ p*. For all conditions in each species, we iterated TPOT over 100 generations to identify the optimized pipeline. We used a population size of 50, which is the number of the best pipelines that are predicted by TPOT in one generation that are then carried to next generation, but with randomly altered parameters in their constituent ML algorithms. We separated 80% of the constructed data sets for training and 20% for validation. The training data subset was used through all steps, while the validation data subsets were kept out of the model during TPOT optimization to prevent data leakage and/or training bias. For each training run, we further applied 10-fold cross-validation to the training data subset. All condition-dependent features were log_10_-transformed before training.

For performance assessment, we employed the adjusted coefficient of determination score 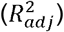. It is often not possible to retrieve feature importance from TPOT-optimized pipelines, since many of the scikit-learn functions selected by TPOT do not support ranking of feature importance. As a result, we assessed how the condition-dependent features impact the predictions by removing them from the data sets and assessing how the TPOT-optimized pipelines of each data set perform. To further assess the prediction results, we used *ϕ* and *E*_*s*_ to obtain predictions about *in vivo* enzyme abundances 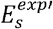 which were compared with the experimental value using the Pearson’s correlation coefficient. We performed GO enrichment analysis using the clusterProfiler R package (Wu et al., 2021; Yu et al., 2012). Further, we performed flux variability analysis (FVA) to assess the variability of central metabolic pathways and of enzyme usage pseudo-reactions.

### Validation of models on genetically modified strains

To assess the validity of CAMEL on unseen conditions and its usefulness for metabolic engineering, we predicted the *in vivo* enzyme abundances 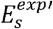 of *E. coli* strains subjected to knock-out mutations followed by adaptive laboratory evolution (ALE) (McCloskey et al., 2018a, 2018b, 2018c, 2018d). For the optimization problem, we replicated the growth conditions by constraining the pcGEM eciML1515 using the estimated 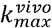 values obtained by Heckmann et al., (2020), which were calculated from quantitative proteomics experiments on the same conditions as McCloskey et al. (2018a, 2018b, 2018c, 2018d). Next, we used the predicted *E*_*s*_ to predict *ϕ* using the machine learning models trained with either the *k*_*cat*_-parameterized dataset or the 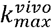 -parameterized dataset. We used the predicted *ϕ* and *E*_*s*_ to obtain calculate the *in vivo* enzyme abundances 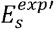 which we then compared with the experimental measurements of Heckmann et al., (2020) using the Pearson’s correlation coefficient.

## Results and Discussion

### Machine learning accurately predicts protein reserve ratios

Our first observation is that the predicted abundance, *E*_*s,i*_, of an enzyme *i* in condition *s* from pcGEMs is usually smaller than the measured *in vivo* abundance 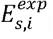. This is the case since predictions on *E*_*s,i*_ must match the corresponding flux, as 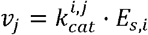, and *E*_*s,i*_ cannot exceed the measured protein abundance. We term this discrepancy between the measured and predicted protein abundance for an enzyme *i* in condition *s* the protein reserve ratio, *ϕ*_*s,i*_, which is calculated by 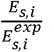.

Since *ϕ*_*s,i*_ cannot be predicted from pcGEMs alone, we rely on machine learning to train models for *ϕ*_*i*_, given data on measured protein abundance and predicted protein usage from pcGEMs over multiple conditions (Figure 1). We refer to the collection **Φ** of reserve ratios over all proteins present in a pcGEM and measured by quantitative proteomic techniques as the distribution of **protein reserve ratios**. Since pcGEMs can be parameterized with *in vitro* measured or with *in vivo* estimated turnover numbers, denoted by *k*_*cat*_ and 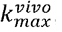, respectively, we performed a comparative analysis of protein reserve ratios in the two parameterization scenarios. Our first observation is that proteins differ with respect to their distributions of reserve ratios, which are dependent on the parameterization of the pcGEMs (Figure 2). For example, the protein reserve ratios of phosphoglycerate kinase in *E. coli* exhibit a narrow distribution for the pcGEM parameterization based on *k*_*cat*_, while the distribution is bimodal for the parameterization with 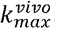. In contrast, the phosphoglycerate kinase in *S. cerevisiae* has the same narrow distribution for both parameterizations. We also observed that the distributions of protein reserve ratios for all investigated enzymes have a heavy tail, due to very large reserve ratios in few conditions.

**Figure 1.**
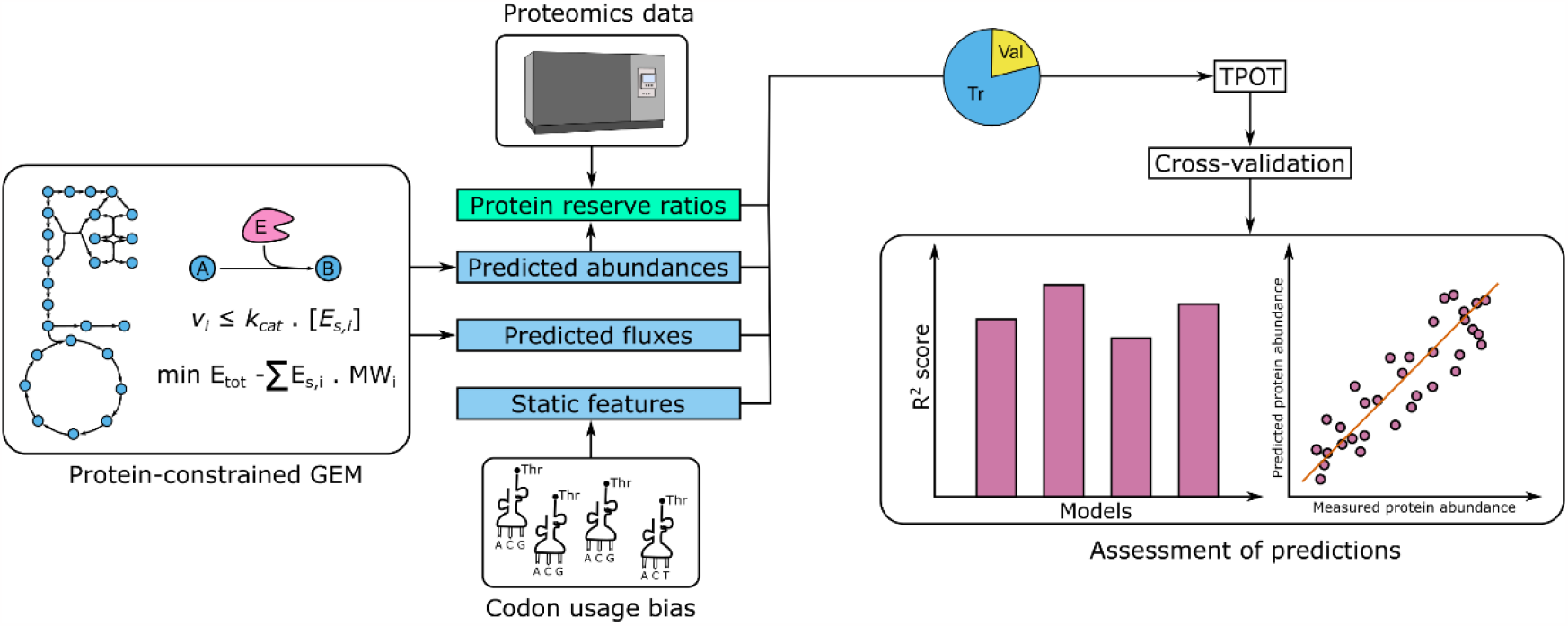
Schematic overview of CAMEL, a constraint-based approach to predict protein abundance. CAMEL uses two types of features, marked as blue boxes: static, obtained from codon usage metrics, and condition-dependent, given by metabolic flux and enzyme usage predicted by constraint-based modelling with pcGEMs. The predicted enzyme usage is employed together with experimental proteomics data to calculate protein reserve ratios that are then used as the target variable for machine learning (green box). The predictive model is obtained by selecting the optimal ensemble of machine learning approaches using TPOT. The performance is assessed by using cross-validation based on regression metrics. Abbreviations: Tr – training; Val – validation.

**Figure 2.**
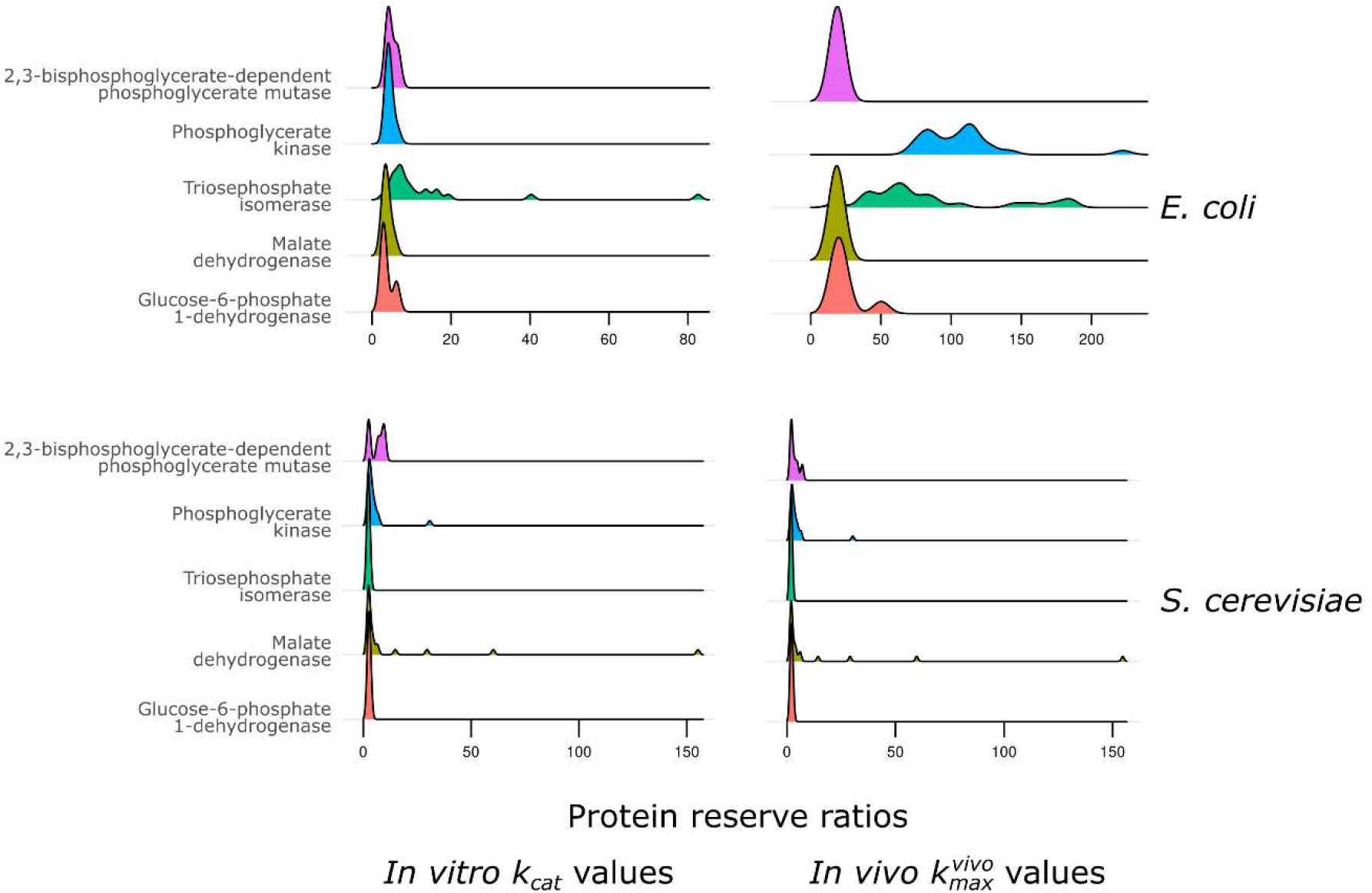
Distribution of protein reserve ratios. The selected proteins take part in central metabolic pathways such as glycolysis, pentose phosphate pathway and citric acid cycle, and are thus present in both organisms. Protein reserve ratios are unitless values.

Next, we used TPOT (Le et al., 2020) to optimize machine learning pipelines to predict **Φ** based on two types of features—static and condition-dependent (Table S1). Static features were obtained from codon usage metrics, while condition-dependent features included enzyme usage and fluxes of the catalyzed reactions predicted by the pcGEM in the two parameterization scenarios (see Material and Methods). Our results indicated that the optimized pipelines for all considered cases exhibited similar structures, composed of two parts: starting with a stacked ensemble of linear and tree-based algorithms and then following with an Extreme Gradient Boosting (XGBoost) algorithm (Chen and Guestrin, 2016), except for the *S. cerevisiae* model using 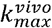 values, whose pipeline included the Extra-trees Regressor (Table S2). Using the trained and optimized pipelines, we then predicted **Φ** for the validation data set, which had not been used during model training and optimization. For *E. coli*, the machine learning models using either the *k*_*cat*_ or 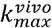 values showed excellent performance 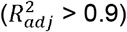, while the model for *S. cerevisiae* showed worse performance 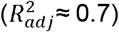 (Figure 3). By comparing the 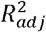 values, we found that for *E. coli*, the pipelines with condition-dependent features derived by using 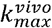 values outperformed those based on *k*_*cat*_ values by 6.7% (Figure 3). For yeast, however, we did not observe a difference in the overall performance of the machine learning models between the two parameterizations.

**Figure 3.**
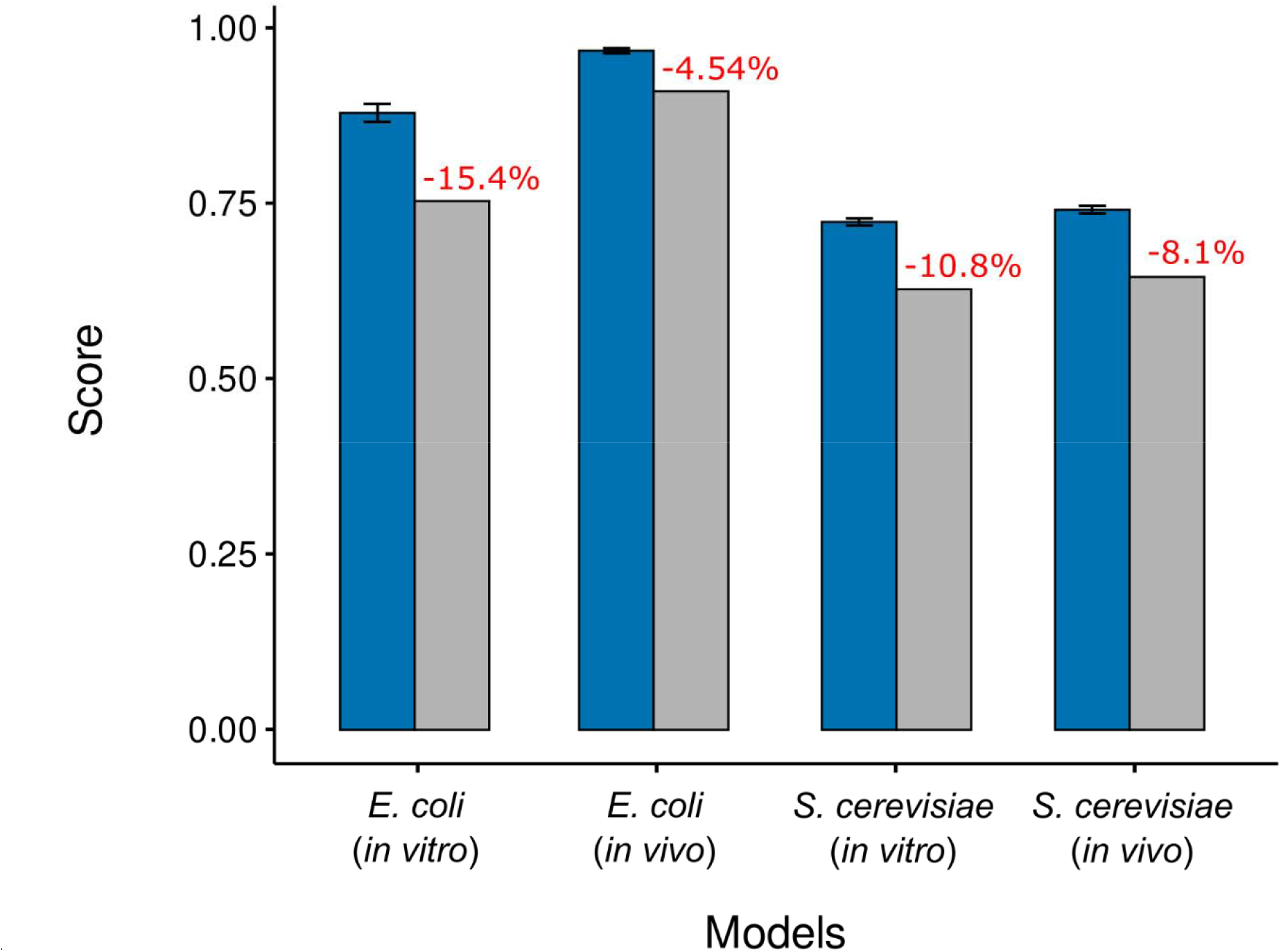
Performance of models from optimized pipelines. The figure shows a comparison of 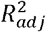 scores for the optimized pipelines based on all features (blue) or excluding condition-dependent features (grey) with either *in vitro k*_*cat*_ or 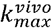 values. Negative percentages (red) indicate a decrease in predictive performance compared to scores obtained from the models trained using all features. Error bars indicate 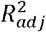 scores obtained from 10-fold cross-validation.

The reasons for the lower predictive performance for yeast in comparison to *E. coli* could be due to several factors: First, from a modelling perspective, there is more and better-curated knowledge about the metabolism of *E. coli* compared to that of *S. cerevisiae* (Bernstein et al., 2023). In addition, the metabolism of yeast is compartmentalized and includes mechanisms for enzyme regulation not accounted for in pcGEMs of any organism (e.g., organellar enzyme pools, post-translational modifications, diffusion effects). Second, from a data perspective, a lack of variability in the protein measurements, resulting from the consideration of samples from similar conditions, can hamper the training of machine learning models. Putting the two perspectives together, the eciML1515 model (1259 enzymes, (Domenzain et al., 2022)) includes a larger number of enzymes in comparison to the ecYeast8 model (965 enzymes, (Lu et al., 2019)), allowing to capture enzymes that may vary more across stress conditions. Lastly, the accuracy of the *k*_*cat*_ values in the BRENDA database (Chang et al., 2021) can also affect the quality of the predictions, given the challenges in measuring them experimentally.

### Importance of fluxes for predicting protein reserve ratios

We were also interested in identifying the extent to which fluxes for reactions associated with a protein affect the predictions of the corresponding protein reserve ratios. To this end, we removed these condition-dependent features from the training data set and re-trained the previously obtained TPOT-optimized pipelines using the reduced training data set. We found that the removal of fluxes resulted in reduced predictive performance in all considered scenarios. For instance, the obtained 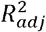 values for *E. coli* were at least 15.4% lower using *k*_*cat*_ values and 4.5% lower using 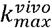. For *S. cerevisiae*, there was a 10.8% decrease for the model using *k*_*cat*_ values and an 8.1% decrease using 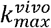 values (Figure 3). These findings demonstrated that the usage of fluxes as condition-dependent features improved model performance in both organisms, particularly in the case when pcGEMs were parametrized with *k*_*cat*_ values. In addition, this shows that the usage of fluxes as features may, to a certain extent, mitigate the effects of model parameterization.

The optimization problem of minimizing excess enzyme usage plays an important role in defining how the flux distribution and enzyme usage distribution are obtained. The objective function ensures efficient utilization of cellular resources, as it avoids burdening the cell with excess enzymes (Bruggeman et al., 2020). We emphasize that, biologically, excess enzymes are important for robustness and adaptability to changing environments, and its minimization could limit the flexibility needed by cells to adapt to these conditions, leading to reduced fitness (Alter et al., 2021). However, excess enzymes result in cellular burden, which can lead to proteotoxic effects for the cell, also reducing fitness (Kintaka et al., 2020). We emphasize that the protein reserve ratios we define do not necessarily correspond to biological protein reserves — we use this terminology to account for differences between predictions from pcGEMs and measured abundances. In that respect, one can use another objective to predict protein fraction used in fluxes; however, the key point is that we will still rely on ratio between measured and predicted values to build ML models — which is the essence of CAMEL.

### Proteins with particular prediction error of protein reserve ratio are enriched in different pathways

Next, we were interested if there are any distinguishing characteristics between the two groups of proteins -- with low or high relative error in predictions. We considered a protein to show low prediction error if the relative error between the predicted and measured protein reserve ratio was lower than 50%, and a high prediction error if the relative error was higher than 100% in each model (noting that varying the definitions with lower, or respectively higher, values have a negligible effect on the conclusions).

First, we checked whether the two groups of proteins were enriched in any specific GO terms (Ashburner et al., 2000; Carbon et al., 2021). For *E. coli*, we found that proteins with low prediction error were enriched in biosynthesis of secondary metabolites, biosynthesis of cofactors, terpenoid biosynthesis and riboflavin metabolism for models using either *k*_*cat*_ and 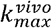 values (Figure S1). For proteins with high prediction error, enriched GO terms shared between models using either *k*_*cat*_ or 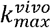 values included biosynthesis of secondary metabolites, carbon metabolism, fatty acid metabolism and degradation (Figure S1). The five proteins with the highest relative error (Table S3) for both model parameterizations were part of the same metabolic pathways, namely: glycolysis, fatty acid biosynthesis, beta-oxidation, and cell wall biogenesis.

Similarly, for *S. cerevisiae*, the proteins with low prediction error for the model using *k*_*cat*_ and 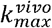 values were enriched in the biosynthesis of secondary metabolites, biosynthesis of amino acids, and carbon metabolism. Likewise, proteins with high prediction error were enriched in biosynthesis of secondary metabolites, biosynthesis of amino acids, carbon metabolism and biosynthesis of cofactors for the models using *k*_*cat*_ and 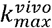 values (Figure S2). The GO term biosynthesis of secondary metabolites encompasses enzymes catalyzing a broad set of reactions, that may explain its significant enrichment in all tested protein sets. The five proteins with the highest relative error (Table S4) showed overlap between the two model parameterizations and were part of purine metabolism, hexose metabolism, and porphyrin-containing compound metabolism. These comparisons demonstrated that the protein reserve ratios did not exhibit distinct patterns based on which metabolic pathway(s) in which the proteins are involved.

To further compare the two groups of proteins, we examined whether the differences in the relative error of predictions were associated with the coefficient of variation (CV) of protein reserve ratios calculated from the models using either *k*_*cat*_ and 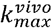 values. For all compared scenarios, many proteins with low CV also had high relative errors. However, we did not identify a statistically significant difference in CV between proteins with low and high relative errors in all scenarios except the *E. coli* model using 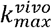 values (Pairwise Wilcoxon rank sum test with Bonferroni correction, Figure S3). Taken together, these results indicated that proteins with low or high prediction errors are used in similar metabolic contexts, suggesting that other physiological factors might contribute to the predictive performance of their abundances.

### CAMEL accurately predicts *in vivo* protein abundances

Next, we used the predicted distributions of protein reserve ratios, 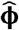, to estimate protein abundance, 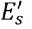, which is given by 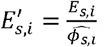, where *E*_*s,i*_ is the condition-specific protein abundance predicted from the pcGEM. The obtained protein abundance estimates were compared to experimental measurements (Chen and Nielsen, 2021; Davidi et al., 2016) by calculating the Pearson correlation coefficient of log_10_-transformed protein abundances for each growth condition separately (shown in Tables S5-S8) as well as the average across all conditions (Figure 4). For *E. coli*, we found a high and significant average value for the Pearson correlation coefficient of 0.987 (p-values < 2.5e-5) over the considered conditions using the models for 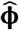 based on the model parameterized with *k*_*cat*_ values, and 0.994 (p-values < 3.1e-11) using the model parameterized with 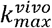 values. In *S. cerevisiae*, using the *k*_*cat*_ parameterization, we found a smaller, yet significant average Pearson correlation of 0.483 (p-values < 0.05 for all significant correlations) over the considered conditions; using the pcGEM with 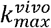 values resulted in an average Pearson’s correlation of 0.513 (p-values < 0.03 for all significant correlations).

**Figure 4.**
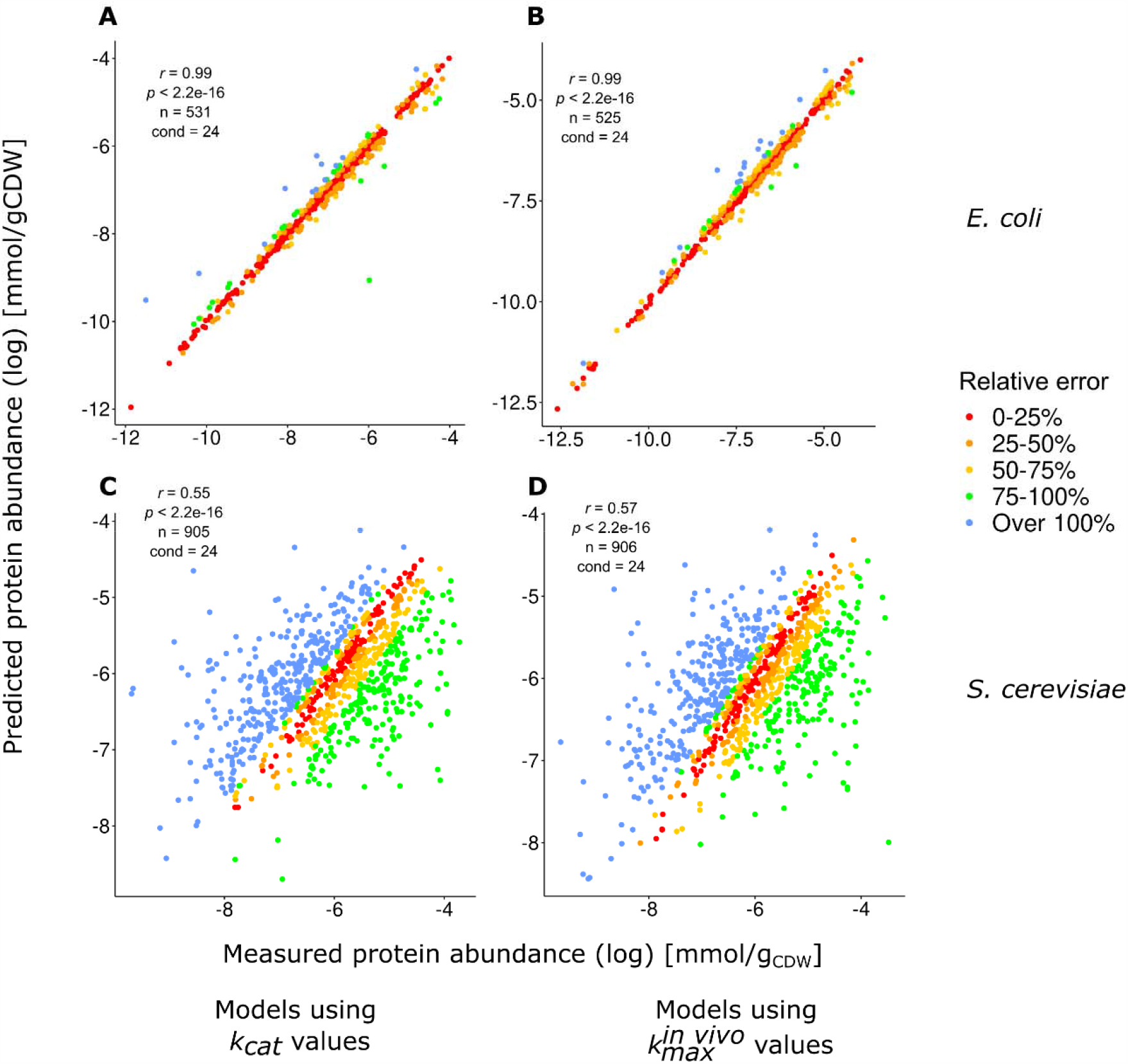
Comparison of measured and predicted protein abundances. The colour scheme indicates the relative error when comparing the measured 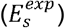 and predicted protein abundances 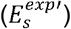. Relative error was calculated using protein abundances in nominal scale. Panel (A) shows predictions using the *E. coli* model with *in vitro k*_*cat*_ values (A), and in vivo 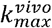 values (B). The panels (C) and (D), depict the prediction performance of the *S. cerevisiae* model with *k*_*cat*_ values (C) and 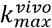 values (D).

Here, too, we took a closer look at proteins with very high relative errors and how their CV compare to proteins with very low relative errors. In *E. coli*, the proteins with the highest relative error showed values larger than 1000% (Table S9). These proteins were part of the chorismate biosynthesis, fatty acid beta-oxidation, pentose phosphate pathway and peptidoglycan biosynthesis when predictions were based on the pcGEM with *k*_*cat*_ values. In the case when 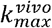 parameterization was employed, these proteins were part of fatty acid biosynthesis, pentose phosphate pathway and purine metabolism. For *S. cerevisiae*, the proteins with the highest errors showed more extreme values than in the case of *E. coli* (Table S10). For both pcGEM parameterizations, the proteins with high error are related to amino acid biosynthesis. As also seen by Lu et al., (2019) and Xia et al., (2022), this observation could be attributed to how different strains and growth conditions affect amino acid biosynthesis pathways. Lu et al. (2019) details that differences in energy pathways used for ATP regeneration directly impacts the biosynthesis of amino acids, given that central metabolic pathways are the main supplier of precursor molecules. Indeed, we observed that some of the proteins with the highest relative error are also related to energy metabolism, such as the pyruvate decarboxylase isozyme 1 and the pyruvate dehydrogenase complex protein X component (Table S10). Xia et al. (2022) have also made a similar observation, stating that the proteome fraction allocated to amino acid biosynthesis is metabolism-dependent. As the cell increases protein translation, the proteome fraction allocated to other sections of metabolism is decreased. However, for the growth conditions we have chosen, the growth rates did not exceed the threshold to trigger the onset of the Crabtree effect. Further, the models were not constrained on amino acid exchange reactions, allowing the model to freely uptake required amino acids. This could explain the high error on these proteins, since they are not being actively used for amino acid biosynthesis. Regarding the CV, we observed no significant difference between proteins with very high or very low errors (Figure S4) (pairwise Wilcoxon rank sum test with Bonferroni correction), except for the yeast pcGEM with 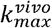 values (p-value < 0.027).

The FVA analysis revealed that for *S. cerevisiae*, there is little variation on metabolic flux through central metabolic pathways (median flux ratio of 16.3 across conditions). For *E. coli*, there was higher variability (median flux ratio of 64.9 across conditions), likely due to the constraints used, since less exchange reactions of central metabolites were constrained than for *S. cerevisiae*. For the enzyme usage pseudo-reactions, we assessed if enzymes with high relative error could be related to pseudo-reactions with high variability. For both *E. coli* and *S. cerevisiae*, using either *k*_*cat*_ or 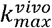 values, there was no relation between the enzyme usage variability and the relative error. Proteins with either high or low relative errors exhibited similar variability. Some proteins with high variability displayed low relative errors, although the difference in enzyme usage variability between the proteins with low variability and proteins with high variability was very small (median ratio variability of 1.0) (Table S11).

To further demonstrate the capabilities of CAMEL, we validated its predictive performance using data from unseen conditions. We predicted the *E*_*s*_ and *ϕ* of mutant *E. coli* strains subjected to ALE. We then used the predictions to calculate the 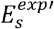 and compared with experimental values using the Pearson correlation coefficient. For the ML trained with the *k*_*cat*_-parameterized pcGEM data, we obtained a value of 0.626 for the Pearson correlation coefficient, while for the ML trained with the 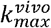 -parameterized pcGEM data, we obtained a value of 0.533 for the Pearson correlation coefficient. This result shows that CAMEL is able to obtain good predictions for genetically modified strains, where the rationale behind metabolic optimality may not hold.

Previous attempts for predicting protein abundance using constraint-based models achieved good predictive performances, but they underestimated *in vivo* protein abundance. The predictions performed by Adadi et al. (2012) using the MOMENT approach achieved a Pearson’s correlation of 0.84 between predicted protein contents and gene expression data of *E. coli* grown on a glucose minimal medium. We showed that CAMEL with 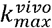 parameterized pcGEM resulted in Pearson correlations of over 0.9 for *E. coli* over different conditions, demonstrating considerably improved performance by using the protein reserve ratios to predict protein usage values compatible to the *in vivo* protein usage. In addition, CAMEL does not require the use of any additional molecular read-outs, thus saving valuable resources for making predictions in unseen conditions. We note that the performance of CAMEL is comparable to that of Resource Balance Analysis (RBA) model of *Bacillus subtilis*, achieving an R^2^ value of 0.94 (Goelzer et al., 2015); however, predictions from RBA models consider more information about transcription, protein translation, translocation, compartmentalization, folding and thermostability, which are difficult to parameterize even in prokaryotic model organisms. CAMEL, on the other hand, depends on pcGEMs, quantitative proteomics data and coding sequences for generating its predictions.

Attempts to predict protein abundance using machine learning have previously achieved good predictive performances but rely on either transcriptomics data or sequence-derived features, which have low correlation to protein content, and are invariable to changes in environmental conditions, respectively (Ferreira et al., 2021; Li et al., 2019). For instance, the model constructed by Terai and Asai (Terai and Asai, 2020), based on three different algorithms, was trained to predict protein abundance in *E. coli* using sequence-derived features and mRNA structure information. The model performance was assessed by Spearman correlation, which ranged from 0.55 to 0.71. Predictions for *S. cerevisiae* were carried out by the Bayesian network constructed by Mehdi et al. (Mehdi et al., 2014), which combined transcriptomics measurements and sequence-derived features, using data from *S. cerevisiae* and *Schizosaccharomyces pombe*. The predictions were evaluated by Spearman correlation, which ranged from 0.61 to 0.77.

Ferreira et al. (Ferreira et al., 2021) generated a predictive model using only codon usage metrics as features, which achieved a Spearman correlation of 0.74. Although the resulting correlations were higher than those from CAMEL, both studies relied only on data obtained from optimal growth conditions. However, CAMEL can be effectively used to make predictions of protein abundance in sub-optimal scenarios, which is a notoriously difficult endeavor (Li et al., 2019). Given that predicted protein abundance based on the protein reserve ratios are in good agreement with experimental values, these results indicated that the coupling of pcGEMs and machine learning can accurately predict the protein allocation *in vivo*.

By coupling constraint-based approaches and machine learning approaches, we demonstrated that our proposed approach, CAMEL, generated accurate predictions of the protein reserve ratios. In addition, we showed that in combination with constraint-based models, CAMEL leverages these machine learning models to obtain a prediction of protein abundance that overcome notable limitations inherent to pcGEMs (e.g., over-accumulation of enzymes, uncertainty in *k*_*cat*_ values). As its most notable merit, CAMEL predicts protein abundance by relying on physiological data, used in the prediction of fluxes, rather than requiring additional omics measurements such as gene expression. Therefore, CAMEL is readily applicable to any condition given that measurements of growth rates and exchange fluxes are available, widening the possibilities for its biotechnological applications (e.g., design of strains). In addition, as our findings demonstrated, CAMEL results in good predictions for protein abundance for sub-optimal conditions, opening the possibility for using these predictions as features to model other complex physiological traits.

## Code and data availability

The code and data can be found in the GitHub repository: https://github.com/mauricioamf/CAMEL

## Author Contributions

Conceptualization, MAMF, PW, MA, WBS and ZN.; Methodology, MAMF, PW, MA, WBS and ZN.; Formal Analysis, MAMF.; Investigation, MAMF.; Writing – Original Draft, MAMF and ZN.; Writing – Review & Editing, MAMF, PW, MA, WBS and ZN.; Supervision, WBS and ZN. Funding Acquisition, WBS and ZN. Project Administration, WBS and ZN.

## Acknowledgments

This study was financed in part by the Coordenação de Aperfeiçoamento de Pessoal de Nível Superior – Brasil (CAPES) – Finance Code 001. P.W. and Z.N. acknowledge the support from the Research Focus Area “Evolutionary Systems Biology” of the University of Potsdam. M.A. and Z.N. acknowledge the support of the Max Planck Society. M.A. and Z.N were supported by the European Union’s Horizon 2020 research and innovation programme, project PlantaSYST (SGA-CSA No. 739582 under FPA No. 664620).

