## Supplementary information for "Accurate prediction of in vivo protein abundances by coupling constraint-based modelling and machine learning"

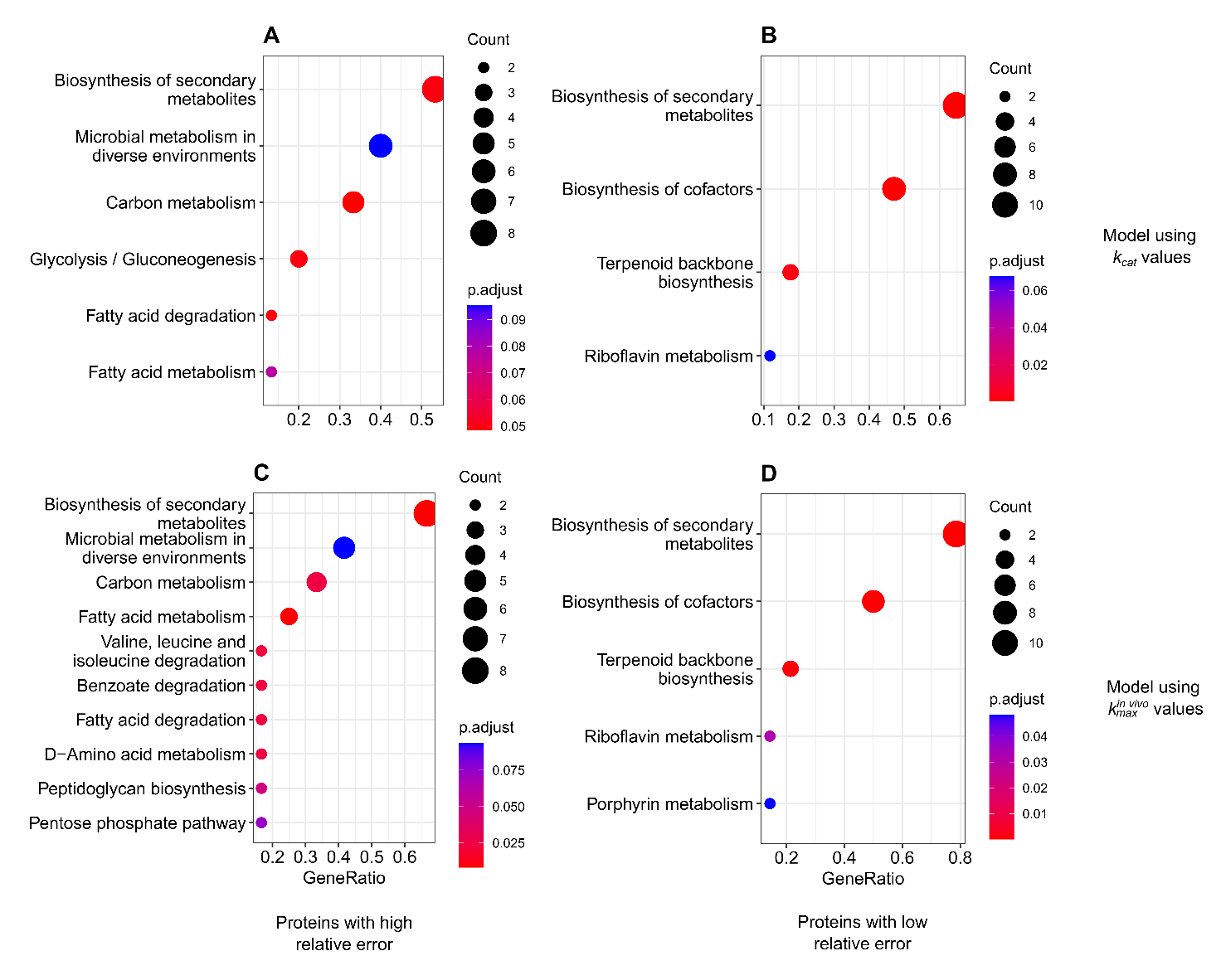

Figure S1. Enriched GO terms for proteins in *E. coli* with high and low relative error for the protein reserve ratio. Count represents the number of proteins assigned the GO term, and GeneRatio denotes the ratio between counts and the sample size. (A) Proteins with high error, model with *in vitro* $\boldsymbol{k}_{\boldsymbol{cat}}$ values, (B) Proteins with low error, model with *in vitro* $\boldsymbol{k}_{\boldsymbol{cat}}$ values (C) Proteins with high error, model with $\boldsymbol{k}_{\boldsymbol{max}}^{\boldsymbol{vivo}}$ values, (D) Proteins with low error, model with $\boldsymbol{k}_{\boldsymbol{max}}^{\boldsymbol{vivo}}$ values.

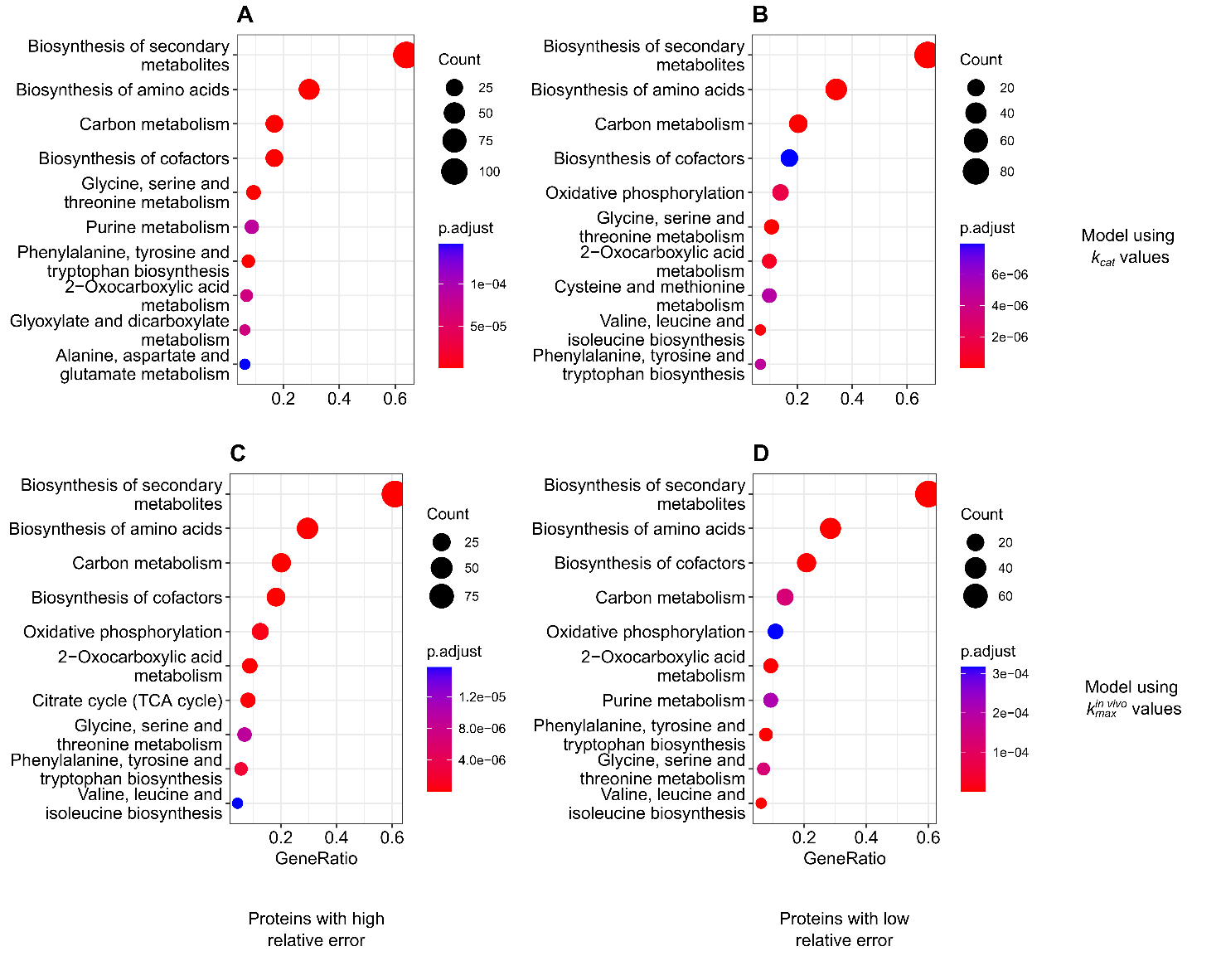

**Figure S2. Enriched GO terms for proteins in *S. cerevisiae* with high and low relative error for the protein reserve ratio.** Count represents the number of proteins assigned the GO term, and GeneRatio denotes the ratio between counts and the sample size. (A) Proteins with high error, model with *in vitro* $k_{cat}$ values, (B) Proteins with low error, model with *in vitro* $k_{cat}$ values (C) Proteins with high error, model with *in vivo* $k_{max}^{vivo}$ values, (D) Proteins with low error, model with *in vivo* $k_{max}^{vivo}$ values.

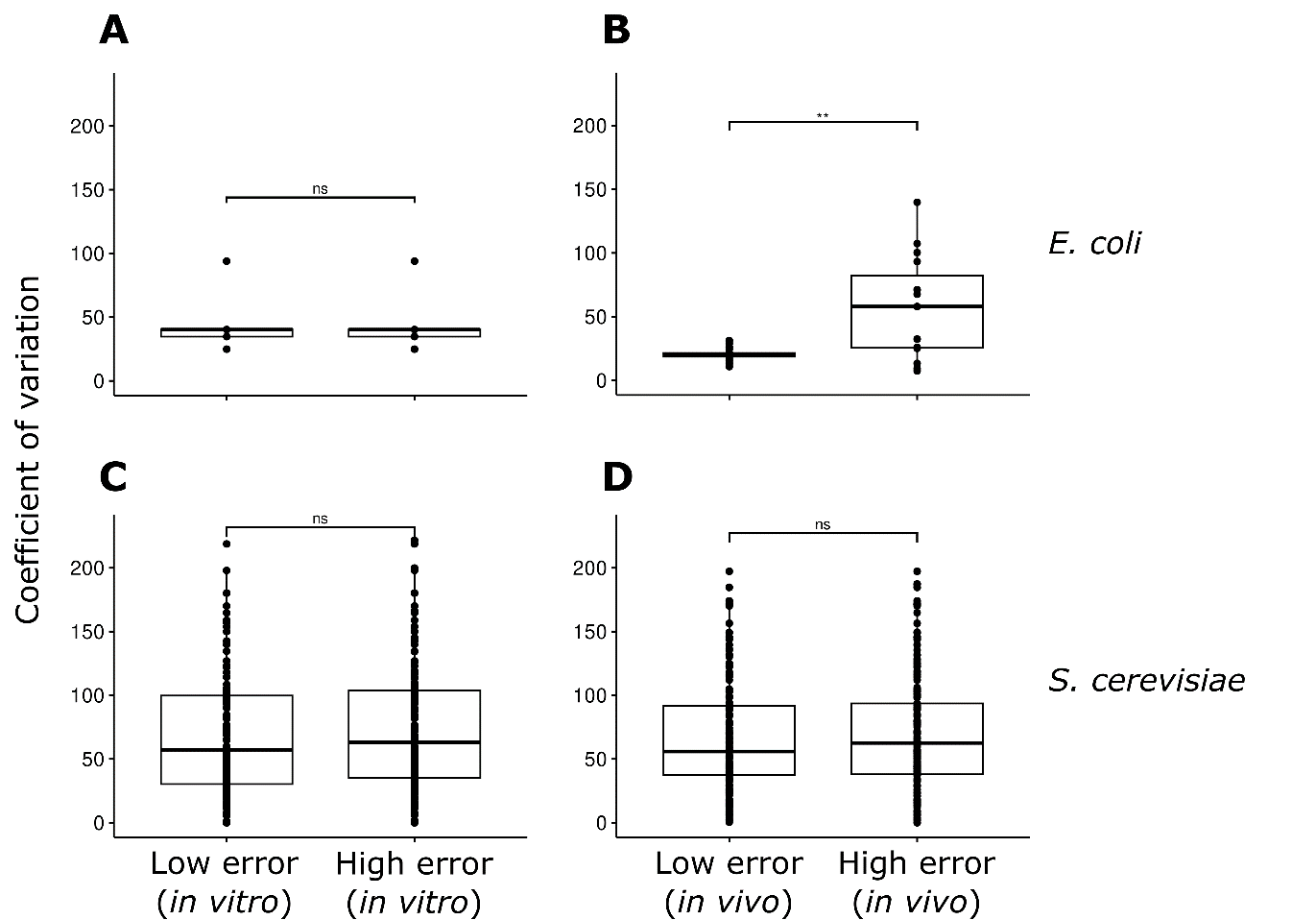

**Figure S3. Comparison of the coefficient of variation between protein reserve ratios with low (≤25%) or high (≥100%) relative errors.** *In vitro* refers to models using *in vitro* $k_{cat}$ values, and *in vivo* refers to models using $k_{max}^{vivo}$ values. (A) Comparison between *E. coli* models using *in vitro* $k_{cat}$ values, (B) *E. coli* models using $k_{max}^{vivo}$ values, (C) *S. cerevisiae* models using *in vitro* $k_{cat}$ values, (D) *S. cerevisiae* models using $k_{max}^{vivo}$ values. A pairwise Wilcoxon rank sum assesses the statistical significance: ** p-value < 0.01.

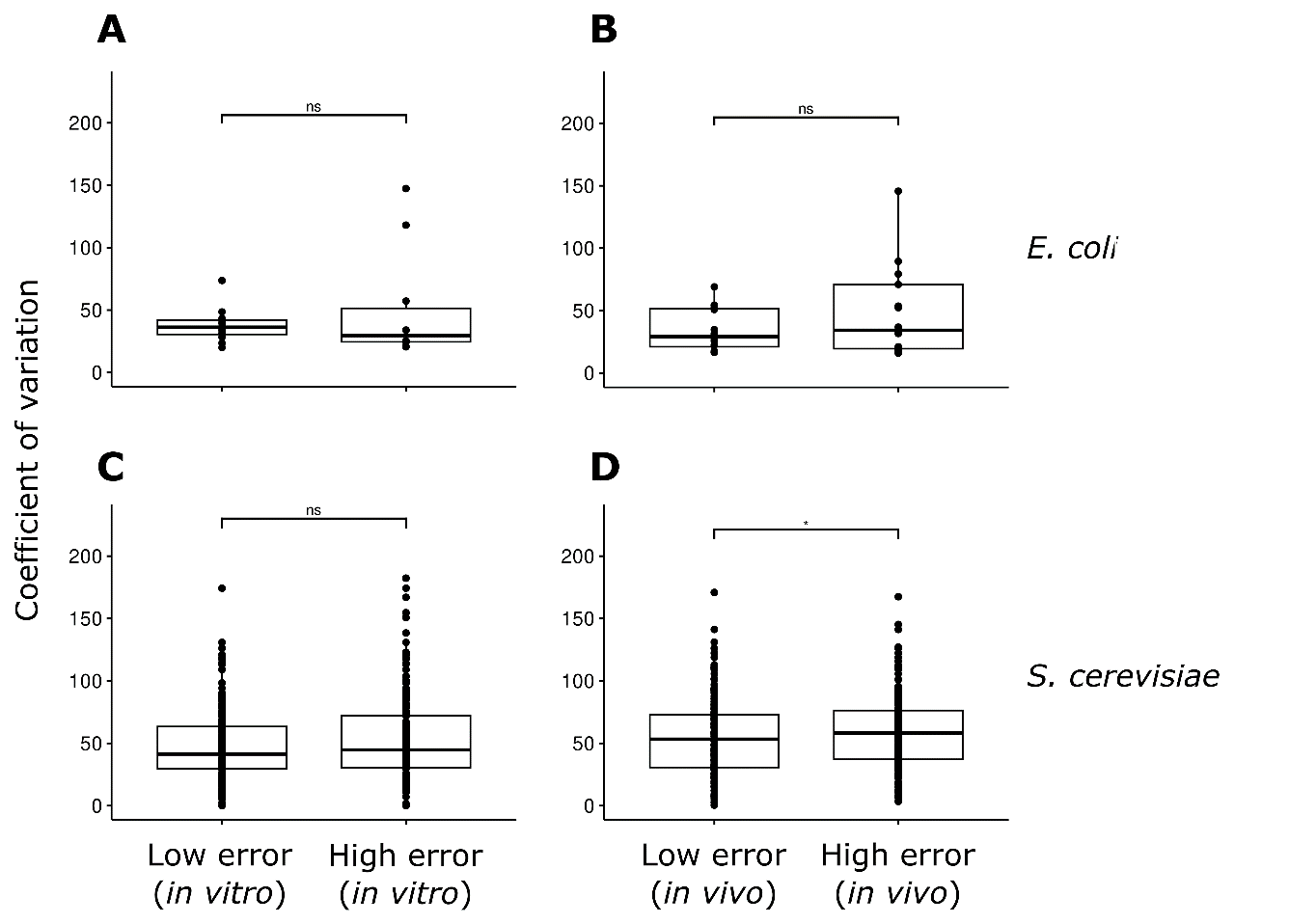

**Figure S4. Comparison of the coefficient of variation between recalculated** $\boldsymbol{E}_{\boldsymbol{s}}^{\boldsymbol{exp'}}$ **values with low (≤25%) or high (≥100%) relative errors.** *In vitro* refers to models using *in vitro* $k_{cat}$ values, and *in vivo* refers to models using $k_{max}^{vivo}$ values. (A) Comparison between *E. coli* models using *in vitro* $k_{cat}$ values, (B) *E. coli* models using $k_{max}^{vivo}$ values, (C) *S. cerevisiae* models using *in vitro* $k_{cat}$ values, (D) *S. cerevisiae* models using $k_{max}^{vivo}$ values. A pairwise Wilcoxon rank sum assesses the statistical significance: * p-value < 0.05.

**Table S1.** Features included in the constructed data sets, for both *E. coli* and *S. cerevisiae*, using predicted $E_{s,i}$ values from models integrated with either $k_{cat}$ or $k_{max}^{vivo}$ values.

| **Features** | **Type** | **Reference** |
| --- | --- | --- |
| Predicted enzyme usage $E_{s,i}$ | Condition-dependent | (Sánchez *et al.*, 2017; Domenzain *et al.*, 2022) |
| Metabolic flux | Condition-dependent | (Savinell and Palsson, 1992a, 1992b) |
| Information theory-based codon usage bias (iCUB) | Static | (Liu *et al.*, 2018) |
| tRNA adaptation index (tAI) | Static | (Reis *et al.*, 2004) |
| Codon adaptation index (CAI) | Static | (Sharp and Li, 1987) |
| Codon bias index (CBI) | Static | (Bennetzens and Hall, 1981) |
| Frequency of optimal codons (Fop) | Static | (Ikemura, 1981) |
| Effective number of codons (ENC) | Static | (Wright, 1990) |
| ENC alternative implementation (ENC’) | Static | (Novembre, 2002) |
| G+C content of gene | Static | (Peden, 2000) |
| G+C of 3^rd^ codon position | Static | (Peden, 2000) |
| Base composition at silent sites | Static | (Peden, 2000) |
| Hydropathicity of protein | Static | (Peden, 2000) |
| Aromaticity of protein | Static | (Peden, 2000) |
| B measure of codon bias | Static | (Karlin *et al.*, 2001) |
| E measure of expression | Static | (Karlin and Mrázek, 2000) |
| Maximum likelihood codon bias (MCB) | Static | (Urrutia and Hurst, 2001) |
| Measure independent of length and composition (MILC) | Static | (Supek and Vlahoviček, 2005) |
| MILC-based expression level predictor (MELP) | Static | (Supek and Vlahoviček, 2005) |
| Synonymous codon usage orderliness (SCUO) | Static | (Wan *et al.*, 2004) |
| Gene codon bias (GCB) | Static | (Merkl, 2003) |
| Evolutionary selection pressure on nucleotide biosynthetic cost (Sc) | Static | (Seward and Kelly, 2018, 2016) |
| Evolutionary selection pressure on gene translation efficiency (St) | Static | (Seward and Kelly, 2018, 2016) |

Table S2. Scikit-Learn (Pedregosa *et al.*, 2011) functions employed in each pipeline optimized by TPOT. Abbreviations: SGDRegressor - Stochastic Gradient Descent regressor; XGBRegressor - eXtreme Gradient Boosting regressor; LassoLarsCV – Cross-validated Lasso using the least-angle regression algorithm.

| ***E. coli* (**$k_{cat}$**)** | ***E. coli* (**$k_{max}^{vivo}$**)** | **Yeast (**$k_{cat}$**)** | **Yeast (**$k_{max}^{vivo}$**)** |
| --- | --- | --- | --- |
| Gradient Boosting Regressor  SGD Regressor  Robust Scaler  XGB Regressor | Function Transformer  Gradient Boosting Regressor  LassoLarsCV  KNeighborsRegressor  XGBRegressor | RidgeCV  Normalizer (L1 norm)  SGDRegressor  XGBRegressor | Function Transformer  PolynomialFeatures  KneighborsRegressor  ExtraTreesRegressor |

Table S3. List of *E. coli* proteins with the highest relative error for the predicted protein reserve ratios, from models using either *in vitro* $\boldsymbol{k}_{\boldsymbol{cat}}$ or $\boldsymbol{k}_{\boldsymbol{max}}^{\boldsymbol{vivo}}$ values.

| **Protein name** | **EC number** | **Data set** |
| --- | --- | --- |
| Triosephosphate isomerase | 5.3.1.1 | $k_{cat}$ |
| Malate dehydrogenase | 1.1.1.37 | $k_{cat}$ |
| 3-ketoacyl-CoA thiolase | 2.3.1.16 | $k_{cat}$ |
| Fatty acid oxidation complex subunit alpha | 4.2.1.17 | $k_{cat}$ |
| Guanyl-specific ribonuclease | 4.6.1.24 | $k_{cat}$ |
| 2,3-bisphosphoglycerate-dependent phosphoglycerate mutase | 5.4.2.11 | $k_{max}^{vivo}$ |
| UDP-N-acetylmuramoylalanine-D-glutamate ligase | 6.3.2.9 | $k_{max}^{vivo}$ |
| UDP-N-acetylmuramate-L-alanine ligase | 6.3.2.8 | $k_{max}^{vivo}$ |
| 3-oxoacyl-[acyl-carrier-protein] synthase 1 | 2.3.1.293 | $k_{max}^{vivo}$ |
| Acyl carrier protein | 2.3.1.40 | $k_{max}^{vivo}$ |

Table S4. List of S*. cerevisiae* proteins with the highest relative error for the predicted protein reserve ratios, from models using either *in vitro* $\boldsymbol{k}_{\boldsymbol{cat}}$ or $\boldsymbol{k}_{\boldsymbol{max}}^{\boldsymbol{vivo}}$ values.

| **Protein name** | **EC number** | **Data set** |
| --- | --- | --- |
| 4-aminobutyrate aminotransferase | 2.6.1.19 | $k_{cat}$ |
| Bifunctional purine biosynthesis protein | 2.1.2.3 | $k_{cat}$ |
| Mitochondrial glycine dehydrogenase | 1.4.4.2 | $k_{cat}$ |
| Phosphoglucomutase 2 | 5.4.2.2 | $k_{cat}$ and $k_{max}^{vivo}$ |
| Delta-aminolevulinic acid dehydratase | 4.2.1.24 | $k_{cat}$ |
| Mitochondrial inorganic pyrophosphatase | 3.6.1.1 | $k_{max}^{vivo}$ |
| Porphobilinogen deaminase | 2.5.1.61 | $k_{max}^{vivo}$ |
| Cytochrome b-c1 complex subunit 8 | 7.1.1.8 | $k_{max}^{vivo}$ |
| Isocitrate dehydrogenase [NADP] | 1.1.1.42 | $k_{max}^{vivo}$ |

Table S5. Pearson correlations between measured ($\boldsymbol{E}_{\boldsymbol{s}}^{\boldsymbol{exp}}$) and predicted ($\boldsymbol{E}_{\boldsymbol{s}}^{\boldsymbol{exp'}}$) protein abundances for each growth condition in *E. coli* using $\boldsymbol{k}_{\boldsymbol{cat}}$ values.

| **Growth condition** | **Pearson correlation** | **p-value** | **Number of proteins** |
| --- | --- | --- | --- |
| ACE_BATCH_mu=0.3_S | 0.997 | 1,74E+02 | 12 |
| GAM_BATCH_mu=0.46_S | 0.891 | 1,90E+04 | 28 |
| GLC_BATCH_mu=0.58_S | 0.998 | 6,69E-18 | 26 |
| GLC_CHEM_mu=0.11_V | 0.979 | 9,29E-11 | 34 |
| GLC_CHEM_mu=0.12_S | 0.979 | 6,88E+00 | 20 |
| GLC_CHEM_mu=0.20_S | 0.997 | 9,28E-18 | 28 |
| GLC_CHEM_mu=0.21_P | 0.993 | 2,88E-18 | 34 |
| GLC_CHEM_mu=0.21_V | 0.995 | 2,73E-14 | 28 |
| GLC_CHEM_mu=0.22_P | 0.998 | 4,28E-31 | 37 |
| GLC_CHEM_mu=0.26_P | 0.986 | 7,40E-08 | 28 |
| GLC_CHEM_mu=0.31_P | 0.988 | 2,58E+09 | 7 |
| GLC_CHEM_mu=0.31_V | 0.988 | 1,20E+06 | 11 |
| GLC_CHEM_mu=0.35_S | 0.998 | 1,67E-05 | 17 |
| GLC_CHEM_mu=0.36_P | 0.995 | 1,04E+00 | 15 |
| GLC_CHEM_mu=0.40_V | 0.998 | 5,24E-14 | 24 |
| GLC_CHEM_mu=0.41_P | 0.995 | 1,97E+00 | 14 |
| GLC_CHEM_mu=0.46_P | 0.998 | 2,54E-06 | 17 |
| GLC_CHEM_mu=0.49_V | 0.954 | 1,44E+08 | 12 |
| GLC_CHEM_mu=0.50_S | 0.992 | 3,07E+01 | 15 |
| GLC_CHEM_mu=0.51_P | 0.997 | 1,62E-09 | 21 |
| GLYC_BATCH_mu=0.47_S | 0.991 | 2,44E-04 | 21 |
| MAN_BATCH_mu=0.47_S | 0.994 | 3,64E-13 | 28 |
| PYR_BATCH_mu=0.4_S | 0.997 | 4,50E-18 | 29 |
| XYL_BATCH_mu=0.55_S | 0.999 | 6,39E-21 | 25 |

Table S6. Pearson correlations between measured ($\boldsymbol{E}_{\boldsymbol{s}}^{\boldsymbol{exp}}$) and predicted ($\boldsymbol{E}_{\boldsymbol{s}}^{\boldsymbol{exp'}}$) protein abundances for each growth condition in *E. coli* using $\boldsymbol{k}_{\boldsymbol{max}}^{\boldsymbol{vivo}}$ values.

| **Growth condition** | **Pearson correlation** | **p-value** | **Number of proteins** |
| --- | --- | --- | --- |
| ACE_BATCH_mu=0.3_S | 0.997 | 3,43E-18 | 28 |
| GAM_BATCH_mu=0.46_S | 0.995 | 4,64E-07 | 20 |
| GLC_BATCH_mu=0.58_S | 0.996 | 1,18E-07 | 21 |
| GLC_CHEM_mu=0.11_V | 0.991 | 5,70E-10 | 26 |
| GLC_CHEM_mu=0.12_S | 0.990 | 2,29E+01 | 16 |
| GLC_CHEM_mu=0.20_S | 0.989 | 3,32E+01 | 16 |
| GLC_CHEM_mu=0.21_P | 0.998 | 1,06E-21 | 29 |
| GLC_CHEM_mu=0.21_V | 0.996 | 3,66E-14 | 26 |
| GLC_CHEM_mu=0.22_P | 0.991 | 4,18E-07 | 24 |
| GLC_CHEM_mu=0.26_P | 0.998 | 2,27E-03 | 14 |
| GLC_CHEM_mu=0.31_P | 0.995 | 5,68E-08 | 22 |
| GLC_CHEM_mu=0.31_V | 0.999 | 2,71E-02 | 11 |
| GLC_CHEM_mu=0.35_S | 0.996 | 3,95E-13 | 26 |
| GLC_CHEM_mu=0.36_P | 0.989 | 2,68E-08 | 27 |
| GLC_CHEM_mu=0.40_V | 0.998 | 3,25E-14 | 22 |
| GLC_CHEM_mu=0.41_P | 0.983 | 1,48E-04 | 25 |
| GLC_CHEM_mu=0.46_P | 0.999 | 7,91E-09 | 17 |
| GLC_CHEM_mu=0.49_V | 0.988 | 3,12E+02 | 14 |
| GLC_CHEM_mu=0.50_S | 0.996 | 5,62E-16 | 27 |
| GLC_CHEM_mu=0.51_P | 0.992 | 7,83E-04 | 20 |
| GLYC_BATCH_mu=0.47_S | 0.998 | 3,64E-34 | 36 |
| MAN_BATCH_mu=0.47_S | 0.995 | 3,53E-05 | 18 |
| PYR_BATCH_mu=0.4_S | 0.997 | 2,68E-11 | 22 |
| XYL_BATCH_mu=0.55_S | 0.994 | 4,01E-03 | 18 |

Table S7. Pearson correlations between measured ($\boldsymbol{E}_{\boldsymbol{s}}^{\boldsymbol{exp}}$) and predicted ($\boldsymbol{E}_{\boldsymbol{s}}^{\boldsymbol{exp'}}$) protein abundances for each growth condition in *S. cerevisiae* using $\boldsymbol{k}_{\boldsymbol{cat}}$ values.

| **Growth condition** | **Pearson correlation** | **p-value** | **Number of proteins** |
| --- | --- | --- | --- |
| Lahtvee2017_EtOH20 | 0.465 | 0.006 | 33 |
| Lahtvee2017_EtOH40 | 0.328 | 0.044 | 38 |
| Lahtvee2017_EtOH60 | 0.295 | 0.080 | 36 |
| Lahtvee2017_Osmo02 | 0.150 | 0.330 | 44 |
| Lahtvee2017_Osmo04 | 0.020 | 0.918 | 27 |
| Lahtvee2017_Osmo06 | 0.442 | 0.031 | 24 |
| Lahtvee2017_REF | 0.074 | 0.652 | 39 |
| Yu2020_Clim | 0.253 | 0.101 | 43 |
| Yu2020_CN115 | 0.717 | 6,06E+06 | 43 |
| Yu2020_CN30 | 0.540 | 0.002 | 29 |
| Yu2020_CN50 | 0.706 | 1,22E+06 | 43 |
| Yu2021_Gln_glc1 | 0.739 | 2,16E+06 | 42 |
| Yu2021_Gln_glc2 | 0.489 | 0.003 | 33 |
| Yu2021_Gln_N30 | 0.559 | 0.0004 | 35 |
| Yu2021_Ile_N30 | 0.785 | 5,18E+05 | 38 |
| Yu2021_Ile_std | 0.697 | 2,02E+07 | 43 |
| Yu2021_N30_005 | 0.535 | 0.0001 | 44 |
| Yu2021_N30_010 | 0.578 | 0.0003 | 34 |
| Yu2021_N30_013 | 0.487 | 0.0005 | 46 |
| Yu2021_N30_018 | 0.336 | 0.0387 | 38 |
| Yu2021_N30_030 | 0.501 | 0.0005 | 44 |
| Yu2021_Phe_N30 | 0.672 | 3,71E+07 | 38 |
| Yu2021_Phe_std | 0.700 | 2,82E+08 | 35 |
| Yu2021_std_010 | 0.531 | 0.0008 | 36 |

Table S8. Pearson correlations between measured ($\boldsymbol{E}_{\boldsymbol{s}}^{\boldsymbol{exp}}$) and predicted ($\boldsymbol{E}_{\boldsymbol{s}}^{\boldsymbol{exp'}}$) protein abundances for each growth condition in *S. cerevisiae* using $\boldsymbol{k}_{\boldsymbol{max}}^{\boldsymbol{vivo}}$ values.

| **Growth condition** | **Pearson correlation** | **p-value** | **Number of proteins** |
| --- | --- | --- | --- |
| Lahtvee2017_EtOH20 | 0.150 | 0.366 | 38 |
| Lahtvee2017_EtOH40 | 0.207 | 0.204 | 39 |
| Lahtvee2017_EtOH60 | 0.430 | 0.012 | 33 |
| Lahtvee2017_Osmo02 | 0.348 | 0.034 | 37 |
| Lahtvee2017_Osmo04 | 0.456 | 0.012 | 29 |
| Lahtvee2017_Osmo06 | -0.107 | 0.539 | 35 |
| Lahtvee2017_REF | 0.434 | 0.023 | 27 |
| Yu2020_Clim | 0.633 | 5,82E+09 | 34 |
| Yu2020_CN115 | 0.591 | 0.0002 | 33 |
| Yu2020_CN30 | 0.730 | 4,00E+06 | 42 |
| Yu2020_CN50 | 0.629 | 2,99E+09 | 37 |
| Yu2021_Gln_glc1 | 0.678 | 5,38E+08 | 36 |
| Yu2021_Gln_glc2 | 0.590 | 4,84E+09 | 41 |
| Yu2021_Gln_N30 | 0.524 | 0.0001 | 48 |
| Yu2021_Ile_N30 | 0.859 | 3,03E+01 | 42 |
| Yu2021_Ile_std | 0.845 | 1,72E+04 | 35 |
| Yu2021_N30_005 | 0.630 | 3,46E+08 | 45 |
| Yu2021_N30_010 | 0.564 | 0.0009 | 31 |
| Yu2021_N30_013 | 0.496 | 0.001 | 40 |
| Yu2021_N30_018 | 0.495 | 0.0006 | 44 |
| Yu2021_N30_030 | 0.443 | 0.002 | 46 |
| Yu2021_Phe_N30 | 0.699 | 5,12E+07 | 40 |
| Yu2021_Phe_std | 0.612 | 0.0001 | 33 |
| Yu2021_std_010 | 0.370 | 0.017 | 41 |

Table S9. List of *E. coli* proteins with the highest relative error for the recalculated $\boldsymbol{E}_{\boldsymbol{s}}^{\boldsymbol{exp'}}$ values, coming from models using either $\boldsymbol{k}_{\boldsymbol{cat}}$ or $\boldsymbol{k}_{\boldsymbol{max}}^{\boldsymbol{vivo}}$ values.

| **Protein name** | **EC number** | **Data set** |
| --- | --- | --- |
| Molybdate-binding protein | 7.3.2.5 | $k_{cat}$ |
| 3-phosphoshikimate 1-carboxyvinyltransferase | 2.5.1.19 | $k_{cat}$ |
| Fatty acid oxidation complex subunit alpha | 4.2.1.17 | $k_{cat}$ |
| 6-phosphogluconate dehydrogenase | 1.1.1.44 | $k_{cat}$ and $k_{max}^{vivo}$ |
| Glutamate racemase | 5.1.1.3 | $k_{cat}$ |
| Flavodoxin 1 | 1.18.1.2 | $k_{max}^{vivo}$ |
| Adenylate kinase | 2.7.4.3 | $k_{max}^{vivo}$ |
| Ribose-phosphate pyrophosphokinase | 2.7.6.1 | $k_{max}^{vivo}$ |
| 3-oxoacyl-[acyl-carrier-protein] synthase 1 | 2.3.1.293 | $k_{max}^{vivo}$ |

Table S10. List of *S. cerevisiae* proteins with the highest relative error for the recalculated $\boldsymbol{E}_{\boldsymbol{s}}^{\boldsymbol{exp'}}$ values, coming from models using either $\boldsymbol{k}_{\boldsymbol{cat}}$ or $\boldsymbol{k}_{\boldsymbol{max}}^{\boldsymbol{vivo}}$ values.

| **Protein name** | **EC number** | **Data set** |
| --- | --- | --- |
| Pyruvate decarboxylase isozyme 1 | 4.1.1.1 | $k_{cat}$ |
| Mitochondrial glycine dehydrogenase | 1.4.4.2 | $k_{cat}$ and $k_{max}^{vivo}$ |
| Asparagine synthetase 1 | 6.3.5.4 | $k_{cat}$ |
| 3-hydroxy-3-methylglutaryl-coenzyme A reductase 2 | 1.1.1.34 | $k_{cat}$ |
| Pyruvate dehydrogenase complex protein X component | 1.2.4.1 | $k_{cat}$ and $k_{max}^{vivo}$ |
| Mitochondrial inorganic pyrophosphatase | 3.6.1.1 | $k_{max}^{vivo}$ |
| NADPH-cytochrome P450 reductase | 1.6.2.4 | $k_{max}^{vivo}$ |
| Ornithine carbamoyltransferase | 2.1.3.3 | $k_{max}^{vivo}$ |

Table S11. Flux variability analysis for central metabolic pathways and enzyme usage pseudo-reactions. Median flux ratio was calculated by dividing the median maximum flux by the median minimum flux across all growth conditions.

| **Species** | **Reactions** | **Dataset** | **Median flux ratio** |
| --- | --- | --- | --- |
| *E. coli* | Central metabolic pathways | $k_{cat}$ | 64,88342 |
|  |  | $k_{max}^{vivo}$ | 52,20936 |
|  | Enzyme usage pseudo-reactions | $k_{cat}$ | 1,000059 |
|  |  | $k_{max}^{vivo}$ | 1,000048 |
| *S. cerevisiae* | Central metabolic pathways | $k_{cat}$ | 16,29839 |
|  |  | $k_{max}^{vivo}$ | 18,62662 |
|  | Enzyme usage pseudo-reactions | $k_{cat}$ | 1,000092 |
|  |  | $k_{max}^{vivo}$ | 1,000089 |
